# Movement-related changes in pallidocortical synchrony differentiate action execution and observation in humans

**DOI:** 10.1101/2020.05.27.117416

**Authors:** Katy A. Cross, Mahsa Malekmohammadi, Jeong Woo Choi, Nader Pouratian

**Affiliations:** Department of Neurology, University of California, Los Angeles; Department of Neurosurgery, University of California, Los Angeles

**Keywords:** Parkinson disease, deep brain stimulation, beta oscillations, action observation, motor control

## Abstract

Suppression of local and network alpha (8-12 Hz) and beta (12-35 Hz) oscillations in the human basal ganglia-thalamocortical (BGTC) circuit is a prominent feature of movement. Local alpha/beta power, cross-region beta phase coupling, and phase-amplitude coupling (PAC) have all been shown to be suppressed during movement in multiple nodes of the BGTC. However, the specificity of these various movement-related changes to actual movement execution remains poorly understood. To differentiate signals that are specifically related to movement execution, we compared changes in globus pallidus internus (GPi) and motor cortical local oscillatory activity and coupling (cross-region phase coupling and local PAC) during movement execution and movement observation in 12 patients with Parkinson disease undergoing deep brain stimulator implantation. We hypothesized that network coupling is more directly related to movement execution than local power changes, given the putative role of pathological network coupling in movement disorders such as Parkinson disease. We observed suppression of alpha/beta power during action observation and execution in the globus pallidus and motor cortex during both action execution and action observation. In contrast, pallidocortical coherence and GPi and motor cortical alpha/beta-gamma PAC were significantly suppressed only during action execution. Our results demonstrate a functional dissociation within the BG-cortical network during action execution and observation in which suppression of BG-cortical functional connectivity and local phase amplitude coupling are features specifically of overt movement, suggesting a particularly important role in motor execution. This has implications for identification and use of intracranial signals for closed loop brain stimulation therapies.

## Introduction

Human movements are associated with changes in oscillatory neural activity across the basal ganglia-thalamocortical motor (BGTC) network. Movement-related suppression of the power of local field potential (LFP) oscillations in the alpha (α, 8-12Hz) and beta (β, 13-35Hz) frequencies and increased power in gamma frequencies (γ, >40 Hz) have been well-established in sensorimotor cortex as well as in the subthalamic nucleus (STN) and globus pallidus internus (GPi) (Pfurtscheller and Lopes da Silva 1999; Miller et al. 2012; van Wijk et al. 2012; Brittain and Brown 2014; Malekmohammadi et al. 2018a). In addition, movement is associated with modulation of phase coupling between alpha and beta signals in the basal ganglia and sensorimotor cortex (Kühn et al. 2006a; Alegre et al. 2010; van Wijk et al. 2017) and cross frequency coupling between alpha/beta phase and gamma amplitude in sensorimotor cortex, STN and GPi (Yanagisawa et al. 2012; de Hemptinne et al. 2013; Tsiokos et al. 2013; Kato et al. 2016; Kondylis et al. 2016; AuYong et al. 2018; Malekmohammadi et al. 2018a). While all these changes are recognized as being movement-related, the functional significance of these distinct signals remains incompletely understood, particularly with respect to which of these signals is critical for and specifically associated with movement execution. Such insights are important to gaining better insight into the pathophysiology of movement disorders such as Parkinson disease (PD) as well as identifying viable and meaningful biomarkers for closed-loop brain stimulation therapies.

Alpha and beta oscillations in the BGTC are often referred to as “anti-kinetic” signals, because they are prominent when the motor system is idling or maintaining a motor state (Engel and Fries 2010; van Wijk et al. 2012; Stolk et al. 2019). The association of suppression in alpha and beta power with movement speed (Gilbertson 2005), force (Stančák et al. 1997), and corticospinal excitability (Sauseng et al. 2009; Mäki and Ilmoniemi 2010) suggest a pivotal role in motor execution. Conversely, movement inhibition (Swann et al. 2009; Solis-Escalante et al. 2012) is associated with increases in beta power and experimentally induced increased beta synchrony in sensorimotor cortex or STN reduces movement velocity (Chen et al. 2007; Pogosyan et al. 2009; Joundi et al. 2012). However, sensorimotor and STN alpha and beta desynchronization are also observed in the absence of overt movement execution, for example during motor imagery (Pfurtscheller and Neuper 1997; Kühn et al. 2006a; Miller et al. 2010; Brinkman et al. 2014), observation of others moving (Gastaut and Bert 1954; Alegre et al. 2005; Hari 2006; Marceglia et al. 2009), and passive movement (Arroyo et al. 1993; Neuper and Pfurtscheller 2001). Thus, while modulation of alpha and beta oscillations in distinct nodes of the BGTC is clearly important in motor processing, suppression of alpha and beta power does not, in itself, indicate movement or even motor preparation. Further work is therefore needed to disentangle the behavioral significance of alpha and beta oscillations.

More recently, studies of neural oscillations have demonstrated that movement modulates not only local power, but also the synchronization or interaction of signals across network nodes. For example, subcortical (STN and GPi) and sensorimotor cortical signals are coupled in the beta band in patients with PD and dystonia (Brown et al. 2001; Cassidy et al. 2002; de Solages et al. 2010; Litvak et al. 2011; Neumann et al. 2015) and this coupling is suppressed during movement (van Wijk et al. 2017; AuYong et al. 2018; Fischer et al. 2019). In addition, phase amplitude coupling (PAC) – a modulation of the amplitude of one frequency by the phase of another – of alpha/beta phase to gamma amplitude in sensorimotor cortex, STN and GPi (Lopez-Azcarate et al. 2010; de Hemptinne et al. 2013; Yang et al. 2014; de Hemptinne et al. 2015; Tsiokos et al. 2017) is dynamically modulated by movement (Miller et al. 2012; Yanagisawa et al. 2012; Kondylis et al. 2016). In contrast to alpha and beta power, only a few studies have examined whether modulation of cross frequency and cross region coupling in the BGTC network occurs in the absence of motor execution, as in the case of motor imagery or action observation. The handful of studies that have assessed this have shown similar desynchronization of STN-cortical alpha/beta coupling during such tasks as during motor execution (Alegre et al. 2005; Kühn et al. 2006a; Marceglia et al. 2009).

The GPi may be important for gating overt movement execution, as it is the primary output nucleus of the motor basal ganglia, which exerts inhibitory control over motor cortex via the thalamus. We therefore hypothesized that, in contrast to the STN, alpha and beta oscillations in the pallidocortical network may uniquely demonstrate a dissociation between motor execution and action observation. In addition, we hypothesized that network-level synchronization may be particularly relevant to movement execution given the exaggeration of cortical-subcortical synchrony in movement disorders, such as PD, and the suppression of this synchrony by therapeutic stimulation (Lopez-Azcarate et al. 2010; Whitmer et al. 2012; de Hemptinne et al. 2015; Oswal et al. 2016; Tsiokos et al. 2017; Malekmohammadi et al. 2018a; Malekmohammadi et al. 2018b). This is in contrast to local oscillatory power in PD, which is sometimes reported as increased at baseline and suppressed with therapy and other times reported as neither abnormally elevated nor modulated by therapy (Kühn et al. 2006b; Weinberger et al. 2006; Ray et al. 2008; Kühn et al. 2009; de Hemptinne et al. 2015; Swann et al. 2015; Weiss et al. 2015). In addition, as recent studies suggest functional dissociations between low (~12-20hz) and high (~20-35hz) beta bands in the BGTC network (van Wijk et al. 2016; Tsiokos et al. 2017), we also examined whether movement specificity is differentially represented in specific sub-bands within the alpha/beta range.

Given well-described activation of the motor system during observation of movements (Fadiga et al. 1995; Hari 2006; Gazzola and Keysers 2009; Kilner and Lemon 2013; Rizzolatti and Sinigaglia 2016), passive action observation provides one way to disentangle signals related to overt movement execution from those related to activation of the motor system in the absence of overt movement. To test these hypotheses, we recorded simultaneous GPi LFPs and sensorimotor cortex electrocorticography (ECoG) during surgical placement of deep brain stimulation electrodes in patients with PD while they executed and passively observed repetitive hand movements.

## Materials and Methods

### Patients

Electrophysiological activity in the right GPi and motor cortex were recorded simultaneously, during periods of alternating rest and action execution and action observation in 10 right-handed patients with idiopathic PD undergoing awake DBS implantation for treatment of PD. 2 additional patients underwent DBS implantation of the bilateral STN and contributed data only from motor cortex ECoG (**Table 1**). Motor cortical ECoG data from 1 GPi patient was not available due to technical error during signal acquisition. In total, analyses described include GPi signals from 10 patients, motor cortex signals from 11 patients, and simultaneous GPi/motor cortex signals from 9 patients. Prior to the studies, all subjects provided written informed consent, as approved by the institutional review board at the University of California, Los Angeles.

**TABLE 1.** Patient demographics.

| **Subject** | **Age** | **Gender** | **Disease duration** | **UPDRS (OFF)** | **Recording Sites\*** |
| --- | --- | --- | --- | --- | --- |
| 1 | 63 | M | 7 | 32 | Motor cortex, GPi |
| 2 | 78 | F | 14 | na | Motor cortex , GPi |
| 3 | 65 | M | 8 | na | Motor cortex , GPi |
| 4 | 66 | M | 9 | 35 | Motor cortex , GPi |
| 5 | 60 | M | 5 | 22 | Motor cortex , GPi |
| 6 | 70 | M | 12 | 40 | Motor cortex , GPi |
| 7 | 71 | M | 6 | 27 | Motor cortex , GPi |
| 8<sup>†</sup> | 80 | M | 16 | 28 | Motor cortex , GPi |
| 9<sup>†</sup> | 61 | M | 5 | 31 | Motor cortex |
| 10<sup>†</sup> | 59 | F | 16 | 71 | Motor cortex |
| 11<sup>†</sup> | 68 | F | 10 | 30 | Motor cortex, GPi |
| 12<sup>†</sup> | 61 | M | 8 | 44 | GPi |

\* right hemisphere, contralateral to side of movement
na = pre-operative UPDRS scores not available
<sup>†</sup> subset of patients who also performed a visual control task

### Behavioral Tasks

All subjects performed each of the execution and observation tasks once, with the order of tasks counterbalanced across participants. Each task involved 30-second blocks of rhythmic left hand opening/closing movements at maximum amplitude with fastest comfortable speed alternating with 30-second rest blocks. Movement/rest periods were cued verbally by the experimenter. In the execution task, patients performed the action with the left hand while wearing a kinematic sensor glove with five piezoelectric sensors that measure finger flexure (5DT data glove 5 Ultra, 5DT Inc., Irvine, CA, USA). In the observation task, patients remained completely still while observing a healthy individual (author NP) perform the same action while wearing the kinematic glove. Subjects were instructed to remain as still as possible while keeping their eyes open during rest and observation blocks and were monitored for compliance by the experimenter. Tasks consisted of 4 to 6 blocks each of rest and movement.

A subset of patients also performed a control observation task in which the stimulus depicted rhythmic visual motion without any human action to begin to explore whether changes associated with action observation were specific to observation of action or related simply to observation of any moving stimulus (see Table 1, 3 GPi recordings, 4 motor cortex recordings). In this visual control task patients passively observed 30-second videos of a ball bouncing at similar frequency to the movements in the observation task alternating with 30-second rest blocks in which they looked at a fixation cross. Four blocks each of ball and fixation conditions were presented. The task was performed in counterbalanced order with observation and execution tasks across subjects.

### Neurophysiologic and kinematic signal acquisition

GPi local field potentials (LFP) were captured via the four ring electrode contacts on the DBS leads at the target position (Model 3387, 1.27 mm lead body diameter, contact length 1.5mm, intercontact distance 1.5mm, Medtronic, Inc., Minneapolis, MN, USA). The DBS lead was targeted to motor (ventral posterolateral) GPi using image-guided targeting, 2-4 mm anterior, 19-24 mm lateral and 4-6 mm inferior to the mid-commissural point (accounting for individual anatomy). All trajectories were confirmed with intraoperative microelectrode recordings of neuronal activity (Israel and Burchiel, 2004) and awake macrostimulation testing of therapeutic and side-effect thresholds at each electrode. Unilateral right sensorimotor electrocorticogram (ECoG) recordings were performed via a non-penetrating subdural strip electrode consisting of eight 4 mm platinum contacts with 1 cm inter-contact spacing (Ad-Tech Medical Instruments, Wisconsin, WI, USA). The ECoG electrode strip was introduced through the same burr hole used for DBS electrode implantation and advanced posteriorly past the central sulcus. Ground and reference electrodes were attached to the scalp. Left hand movements synchronized to neural recordings were transduced via the kinematic sensor glove worn by the patient (movement condition) or experimenter (observation condition). Neurophysiologic and glove signal acquisition was performed using BCI2000 v2 or v3 connected to a standalone amplifier (g.Tec, g.USBamp 2.0) with a sampling rate of 2400 Hz following 0.1 Hz-1000 Hz online band-pass filter.

### Anatomical localization of DBS leads and ECoG electrode strip

Confirmation of anatomical localization of the GPi DBS electrode contacts was performed using Lead DBS software for 7 of 9 subjects with available post-operative imaging (Horn et al. 2019). Postoperative CT scans were co-registered to preoperative T1-weightd structural MRI (MPRAGE, slice thickenss 1mm, 3T Siemens Skyra) with two-stage linear registration (rigid followed by affine) using the SyN registration approach as implemented in advanced normalization tools (ANTs) (Avants et al. 2008). Automated reconstruction of electrode trajectory and contact locations was performed using the PaCER toolbox (Husch et al. 2018). Electrode locations were then confirmed on the high resolution structural MRI in patient native space. The most ventral pair of electrodes were verified to be located within the GPi and bipolar LFP recordings from these 2 most ventral electrodes was used for all analyses.

Anatomical localization of the ECoG electrode contacts was performed using 2D/3D fusion techniques adapted from Randazzo et al. (Randazzo et al. 2016) by registering the intra-operative fluoroscopic image to the reconstructed cortical surface of the pre-operative MRI. The pair of electrodes immediately anterior to the central sulcus was identified as motor cortex and bipolar ECoG signal from this pair were used for subsequent analyses (Figure 1A). For two patients localization was not possible due to absence of postoperative imaging or intraoperative fluoroscopy data; for these patients contacts with greatest beta modulation during movement were selected as a proxy for primary motor cortex (Hari and Salmelin 1997).

**Figure 1.**
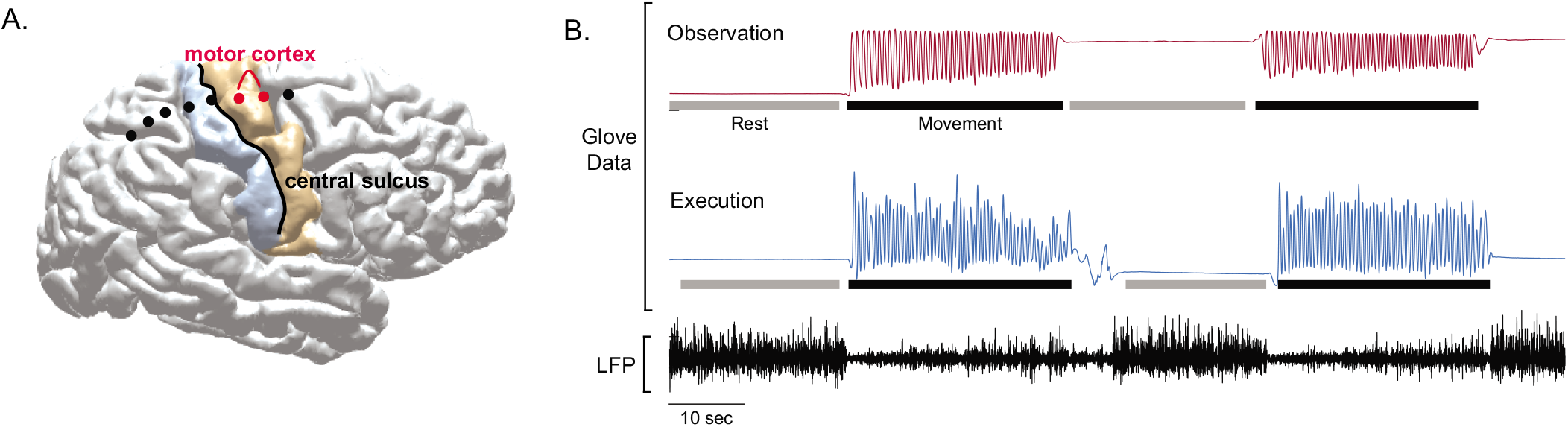
(A) Example of individual subject cortical reconstruction with ECoG electrode localization. Motor cortex recordings were obtained from bipolar electrode pair immediately anterior to the central sulcus. (B) Example glove data (top) and LFP recording from motor cortex during execution block. Bars below glove data indicate periods selected for rest (gray) and movement (black) periods. Movement was defined by onset and offset of clear rhythmic finger flexions. Rest periods were defined by absence of movements with brief (~1s) padding before and after rhythmic movement onset to account for transition period during which patient lifted arm from resting position.

### Neurophysiologic Signal Processing

Right motor ECoG and right GPi LFPs were analyzed along with concurrent left hand kinematic data in Matlab (Mathworks Inc., Natick, MA) utilizing the Fieldtrip toolbox (Oostenveld et al. 2011) and custom scripts. Raw ECoG and LFPs were lowpass filtered at 400 Hz and highpass filtered at 1 Hz using onepass zero-phase FIR filter (lowpass transition band width 100, highpass transition band width 2) (Widmann et al. 2015). Line noise at 60 Hz and harmonics were filtered out with bandstop Butterworth filter (order 4). Bipolar re-referencing between adjacent electrodes was then performed to emphasize local voltage changes and minimize common noise. By careful visual inspection of the movement traces recorded by the data glove, electrophysiological data were segmented into rest and movement blocks (Figure 1B). LFP data from selected blocks were then visually inspected for electrical artifacts (sudden large amplitude shift) while blinded to condition. Blocks with at least 16 seconds of contiguous artifact-free data were included in analyses. Rest and movement block lengths were then matched for each task condition by truncating longer blocks to remove any difference in timepoints from contributing to condition differences. This resulted in an average of 115 seconds per condition (range 65-154).

Preprocessed LFP and ECoG data were decomposed into their time-frequency representation by convolution with a set of complex morlet wavelets. The wavelet family was defined as a set of Gaussian-windowed complex sine waves at 100 logarithmically spaced frequencies between 2 Hz and 300 Hz. For power analyses, wavelet width was 10 cycles for frequencies under 40 Hz and 25 cycles for frequencies above 40 Hz to minimize frequency smoothing and allow for identification of sub-bands within the alpha/beta range. For phase coupling and cross frequency coupling measures requiring instantaneous phase and amplitude values, wavelet width was increased from 3 to 15 cycles in logarithmically spaced steps to emphasize temporal precision (Cohen 2014). The resulting complex analytic signals provided the input for subsequent power, phase coherence and phase amplitude coupling analyses.

Power spectra for each of the 4 conditions (Execution/Observation x Movement/Rest) within each subject were obtained by taking the squared complex magnitude of the analytic signal and averaging over time within each block, and subsequently over blocks within each of the 4 conditions for each subject. To minimize the interregional and inter-subject baseline power differences, for each task the resulting raw power spectra were normalized by dividing each frequency by the total power at rest between 6 and 200 Hz (excluding line noise and harmonics) and converted to decibel scale.

Phase based connectivity between GPi and motor cortex was evaluated using the phase locking value (PLV) (Lachaux et al. 1999), which is defined as the magnitude of the mean phase difference between the two signals expressed as a complex unit-length vector according to the formula 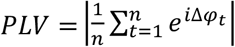 where Δ*φ_t_* is the phase difference between the two signals at time t. If the signals’ phases are completely independent, the relative phase will have a uniform distribution over time and the PLV is zero. Conversely, if the phases of the two signals are strongly coupled then the relative phase (i.e. phase difference) will be clustered and PLV will approach one. PLV was chosen as the measure of interregional coupling, because in contrast to cross-spectral coherence, changes in PLV are insensitive to fluctuations in power that are known to be present in the conditions of interest. Similar to power analyses, PLV was calculated for each frequency in each 30-second block, and subsequently averaged over blocks within each of the 4 conditions for each subject.

Finally, local cross frequency phase-amplitude coupling in GPi and motor cortex was calculated using the debiased Phase-Amplitude Coupling (dPAC) method (Lin et al. 2006; van Driel et al. 2015), which is comparable to other commonly used methods but may have higher sensitivity in the presence of noise (van Driel et al. 2015). dPAC describes interactions between activity at different frequencies in which the amplitude of a signal at one frequency is modulated by the phase of a lower frequency signal. In BGTC network, coupling between alpha/beta phase and gamma amplitude is modulated by movement (Yanagisawa et al. 2012; de Hemptinne et al. 2013; Kato et al. 2016; Kondylis et al. 2016; Tsiokos et al. 2017; AuYong et al. 2018) and we therefore focused on these frequency bands. To calculate the dPAC, instantaneous phase angles were extracted as the angle of the analytic signal at the frequencies of interest for phase (4-40Hz) and corresponding instantaneous amplitudes were extracted as the squared absolute value of the analytic signal at the frequencies of interest for amplitude (30-300 Hz). Vectors in polar space were then defined at each timepoint by the angle of the frequency for phase and the length by the power of the amplitude-modulated frequency. To reduce any potential bias introduced by non-uniformity in the phase angles of frequency for phase, a debiasing term is introduced by subtracting the average vector of the modulating phase angles (Van Driel, 2015). The length of the average of power-adjusted phase angle vectors over time provides the dPAC, which is mathematically defined as 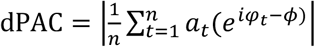 where 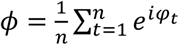, *n* is the number of time points, *a_t_* the amplitude of the modulated frequency at time *t* and *φ_t_* is the phase of the modulating frequency at time *t*. dPAC is zero if there is no relationship between power and phase and is greater than zero when there is a relationship between power and phase. To normalize across frequencies and patients, dPAC values were transformed into z-values at the single subject level by comparing against surrogate values from 1000 permutations in which the power values were shuffled relative to the phase values (Canolty et al. 2006; Cohen 2014).

### Group Statistical Analyses

We first aimed to define which frequency bands were significantly modulated by movement during observation and execution for each neurophysiological measure (normalized power, PLV, dPACz) across the entire frequency space. Movement and rest blocks were compared within each recording site (GPi, M1) and task (execution, observation) using permutation based non-parametric paired tests at the group level with cluster correction for multiple comparisons across frequencies. Implementation of paired tests between movement and rest blocks was accomplished by creating a null distribution of the condition difference by swapping the sign of a subset of pairwise differences in all possible permutations (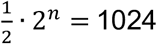 permutations for M1, 512 permutations for GPi)(Nichols and Holmes 2002). P-values were then defined by the probability of the observed condition difference based on the permuted null distribution, with a two-tailed probability of less than 5% considered significant. To correct for multiple comparisons across frequencies, a null distribution of maximum cluster size (number of significant neighboring frequencies based on two-tailed p<0.05) was obtained based on the same permutations and only those observed cluster sizes with a probably of less than 5% as compared to the null distribution were considered significant (Nichols and Holmes 2002; Maris and Oostenveld 2007).

Next, to determine whether differences between tasks and frequency bands were significant, we performed repeated measures ANOVA for each neurophysiologic measure using the magnitude of movement modulation (Movement – Rest difference score) averaged over selected frequency bands as the dependent variable and factors frequency band and task (Execution, Observation) as repeated measures. We were interested in potentially dissociable frequency bands within the broader alpha/beta range (8-35 Hz) known to covary with execution and observation tasks. However, as frequency cutoffs used in the motor neurophysiology and PD literatures are variable (common examples are 8-14 Hz and 15-25 Hz in motor physiology and 12-20 Hz and 20-35 Hz in PD), we used a data-driven approach to select relevant sub-bands within this range for use in the ANOVA similar to previous studies in PD that have identified dissociable “low beta” and “high beta” bands (van Wijk et al. 2016; Tsiokos et al. 2017). Peaks were identified in the power spectra for each subject, task condition and region (9 patients x 4 conditions x 2 regions and 3 patients x 4 conditions x 1 region = 84 spectra) between 8 and 35hz using matlab “peakfind.m” function and confirmed with visual inspection. This was performed across all conditions and brain regions to avoid biasing subsequent statistics. A histogram of the identified peaks showed a bimodal distribution (Figure 2) separating the window from 8-35 Hz into two bands, alpha/low beta (α/low-β) and high beta (high-β), without a clear distinction between alpha and low beta. To identify the specific cut-off frequency between these two bands, a mixture of two Gaussian distributions was fitted using the Matlab curve fitting toolbox and the lowest point between the two distributions used as the boundary between two distinct frequency bands centered at 13.0 Hz (α/low-β) and 23.4 Hz (high-β) with a cutoff between bands at 17.9 Hz. Examination of the Akaike information criterion values confirmed that the bimodal mixture model was superior to either single Gaussian distribution or a mixture of 3 Gaussian distributions. For dPAC analyses, gamma frequencies in the amplitude modulated signal were separated into low gamma (50-90 Hz), high gamma (90-200 Hz,) and high frequency oscillations (HFO, 200-300 Hz), motivated by prior studies demonstrating movement-modulated PAC in PD between beta and low gamma/HFO frequencies in the GPi (Tsiokos et al. 2017; AuYong et al. 2018) and between beta and broadband high gamma in motor cortex (de Hemptinne et al. 2013; Kondylis et al. 2016).

**Figure 2.**
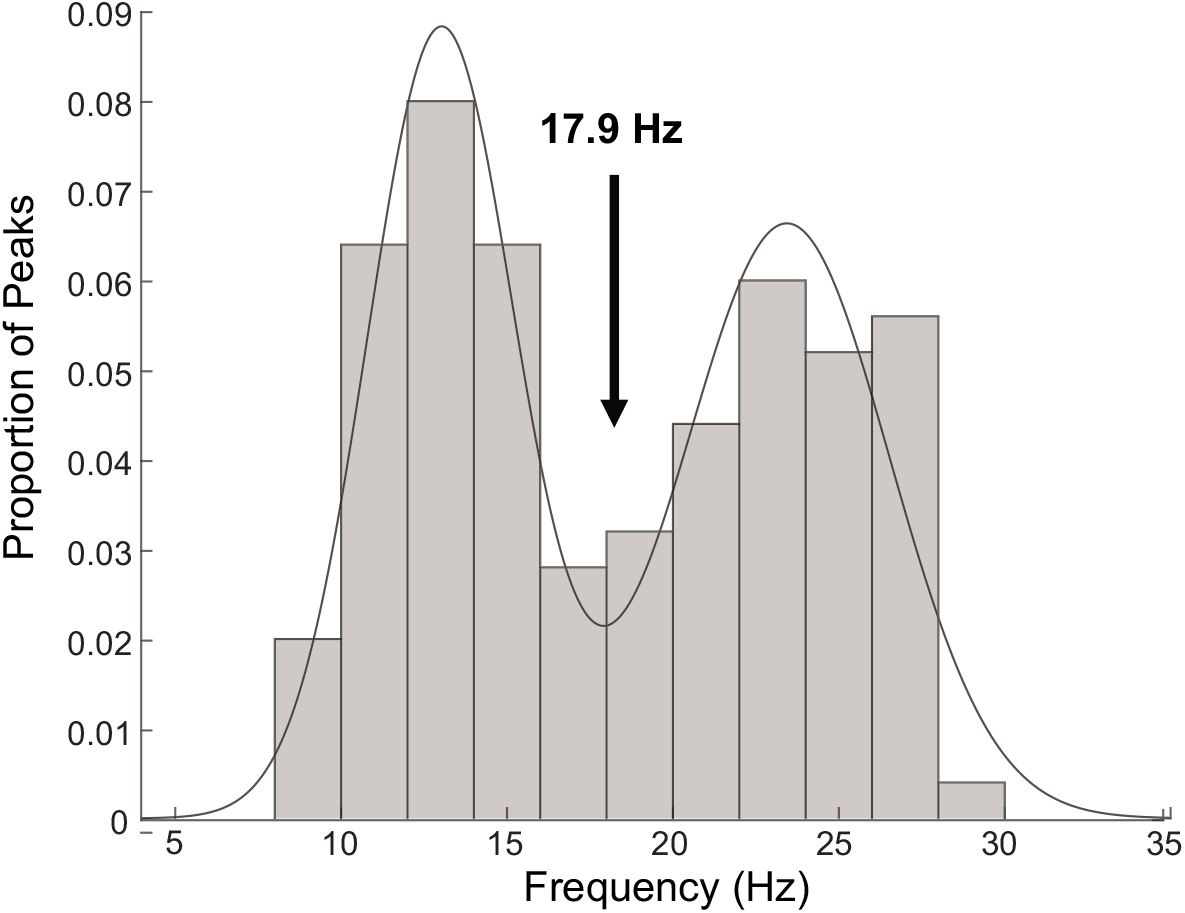
Histogram of individual subject peak frequencies within alpha/beta range (8-35 Hz). Line shows mixture of 2 gaussian models with minimim defining border between identified sub-bands.

Separate ANOVAs were performed for each measure (normalized power, PLV and dPACz) and brain region (motor cortex, GPi). For power and PLV, task (execution, observation) and frequency band (α/low-β and high-β) were included as two repeated measures factors. For dPAC three repeated measures factors were task (execution, observation), phase encoding frequency band (α/low-β and high-β) and amplitude modulated frequency band (low γ, high γ and HFO). Post-hoc contrasts investigating significant effects were carried out (STATA *contrasts* command) and p-values corrected for multiple comparisons using Bonferroni method (reported as p_corr_). In addition, planned comparisons to determine significant movement suppression within each data-driven frequency band was also assessed by comparing the magnitude of movement modulation to zero using one-sample Wilcoxon signed-rank tests (one-tailed given directional hypothesis) with Bonferroni correction for multiple comparisons. These analyses were implemented in Stata 14 software (StataCorp LLC, College Station, TX).

## Results

### Power is suppressed during action execution and action observation

Comparisons of spectral power between rest and movement during execution and observation are shown in Figure 3. Patterns of power modulation were similar in the motor cortex and GPi with movement-related suppression of alpha and beta power during both execution and observation. Non-parametric permutation tests across the frequency space demonstrate that whereas execution led to suppression of a broad band of alpha and beta frequencies (5-35 Hz) in both GPi and motor cortex, observation was associated with significant suppression of a narrower band (8-20 Hz) including alpha and low beta, but not high beta frequencies, in both regions. Results were equivalent when comparing each task’s movement block to a single rest power spectrum (averaged across tasks), confirming that observed differences were due to differences in movement modulation and not rest period activity.

**Figure 3.**
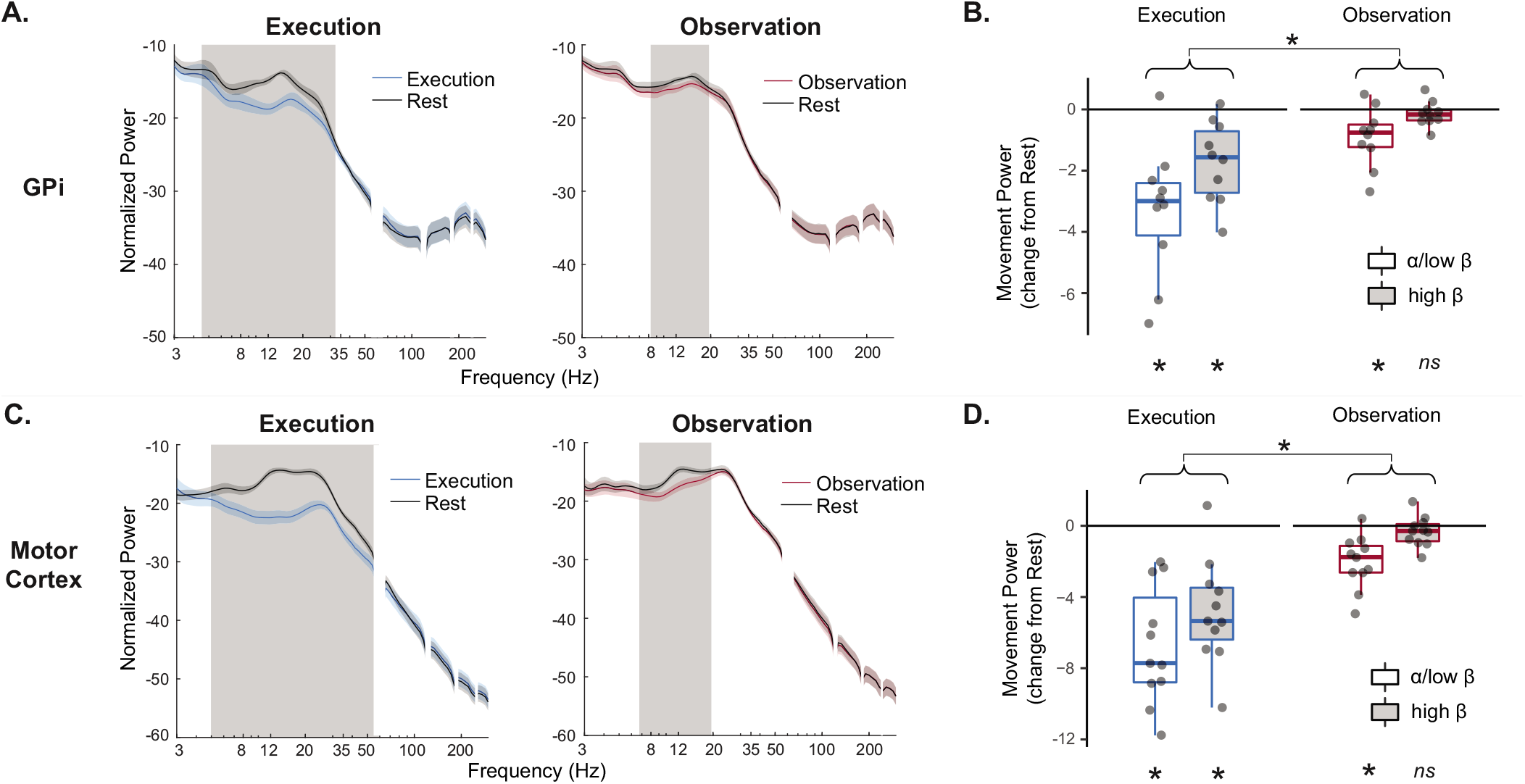
Spectral Power. Normalized spectral power in GPi (A) and motor cortex (C) across frequencies for movement and rest blocks during execution (blue) and observation (red). Gray shading indicates frequencies in which movement and rest are significantly different by non-parametric permutation testing with cluster correction for multiple comparisons across frequencies. Boxplots show power modulation in GPi (B) and motor cortex (D) during movement relative to rest (movement – rest change score) during execution (blue) and observation (red). Significant modulation by movement (e.g. significant difference from zero by Wilcoxon signed rank tests) is indicated at bottom. Main effect of task is indicated at top; main effect of frequency band was also significant but not shown for readability. Given the absence of a two-way interaction, simple effects were not tested. (* p<0.05 Bonferroni corrected), ns non-significant)

Similar effects were demonstrated in statistical tests of band averaged power (Figure 3B/D). In motor cortex, significant movement-related suppression (i.e. movement modulation < 0 by one-sample Wilcoxon signed rank test) was observed during execution in both α/low-β (z=2.93, p_corr_=0.002) and high-β (z=2.845, p_corr_=0.004) whereas during observation only α/low-β (z=2.845, p_corr_=0.004) was significantly suppressed (high-β, z=1.33, p_corr_=0.41). The same pattern was observed in the GPi with significant movement-related suppression in all frequency bands except high-β (Execution: α/low-β z=2.70, p_corr_=0.008; high-β z=2.70, p_corr_=0.008. Observation: α/low-β z=2.40, p_corr_=0.03; high-β z=1.27, p_corr_=0.46).

2×2 repeated measures ANOVA directly comparing the effects of frequency band (α/low-β, high-β) and task (execution, observation) on movement modulation were significant in both regions (GPi F_12_=5.0, p<0.001; M1 F_13_=6.39, p<0.001). Similar patterns were observed in GPi and M1 with significant main effects of frequency band (GPi F_1,9_=10.1, p=0.004; M1 F_1,10_=8.2, p=0.008) and task (GPI F_1,9_= 28.6, p<0.001; M1 F_1,10_=53.2, p<0.001), reflecting greater power suppression for execution compared to observation (GPi t=3.18, p=0.004; M1 t=7.29, p<0.001) and for α/low-β compared to high-β (GPi t=5.34, p<0.001; M1 t=2.86, p=0.008). The interaction between frequency band and task was not significant, indicating that the absence of modulation of high-β during observation may have resulted from the overall smaller magnitude of power changes during observation as compared to execution.

Qualitative comparison of power during action observation and ball observation in the subset of patients who performed the visual control task (n=3 GPi, n=4 M1) provides preliminary evidence that power suppression in GPi may be specific to observation of human actions, as previously described in motor cortex and STN (Hari et al. 1998; Marceglia et al. 2009; Alegre et al. 2010). Suppression of alpha/beta power during observation of a moving ball was less pronounced than that observed during action observation in these subjects (Figure 4).

**Figure 4.**
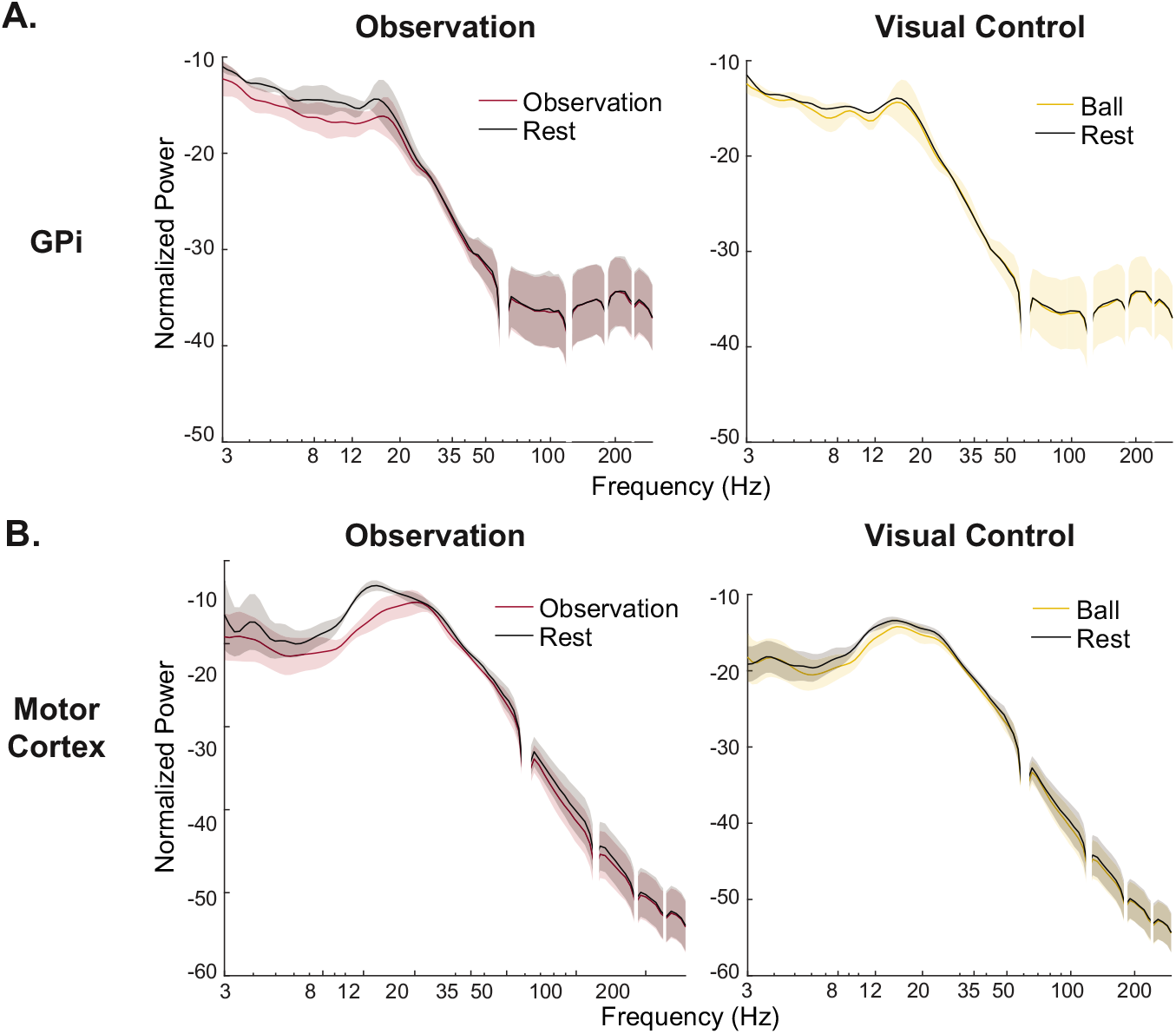
Visual Control Task. Normalized spectral power in GPi (A) and motor cortex (B) across frequencies for movement and rest blocks during observation (red) and ball observation (visual control, yellow) in subset of patients performing both tasks show greater suppression during action observation than ball observation.

### GPi-M1 phase coupling is suppressed during action execution but not action observation

Significant suppression in phase coupling (PLV) between GPi and M1 was observed in α/low-β (9-14 Hz) during execution. In contrast, observation showed no significant suppression of pallidocortical phase coupling (Figure 5A). Again, there were no significant differences between rest conditions in the two tasks and results were equivalent when comparing each task’s movement block to the same rest data obtained by averaging rest periods across tasks.

**Figure 5.**
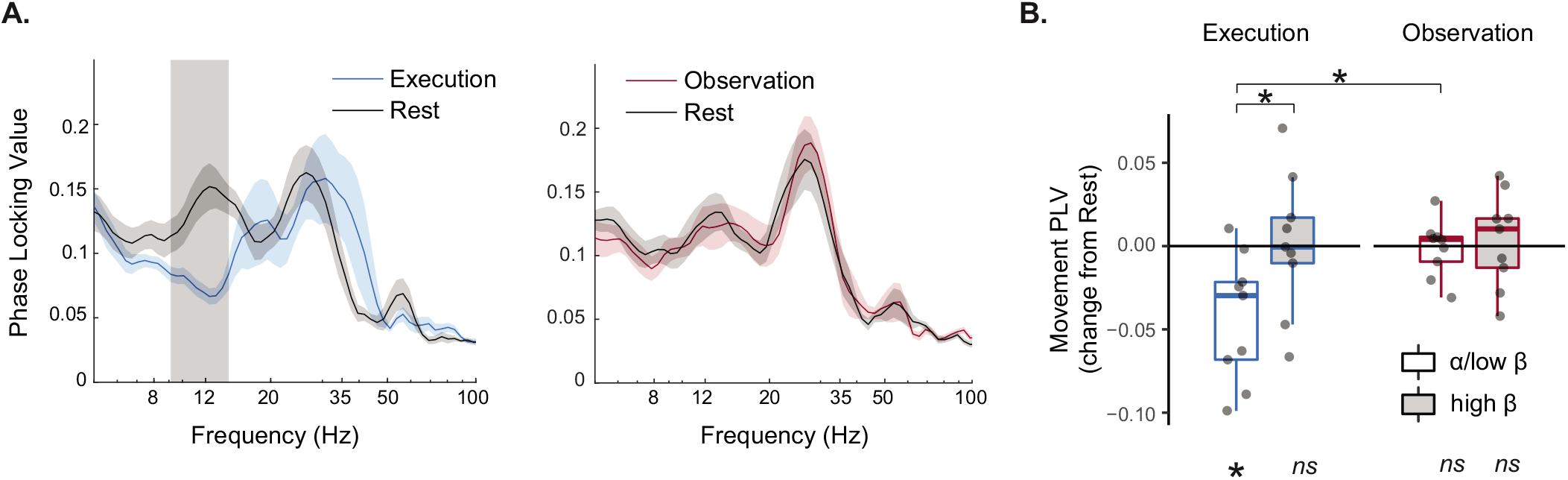
GPi-M1 phase coupling (PLV). (A) PLV across frequencies for execution and observation tasks shows significant movement-modulated PLV only during execution limited to the α/low-β band. Gray shading indicates frequencies in which movement and rest are significantly different by non-parametric permutation testing with cluster correction for multiple comparisons across frequencies. (B) Boxplot shows GPi-M1 PLV modulation during movement relative to rest (movement – rest change score) during execution (blue) and observation (red). Significant modulation by movement (e.g. significant difference from zero) is shown at bottom, and is present only in the α/low-β band during execution. Paired contrasts from significant task x frequency band interaction are shown at top. (* p<0.05 bonforroni corrected; ns = non-significant)

Planned comparisons of band extracted PLV corroborate these results, with significant modulation of phase coupling present only in the α/low-β band during movement execution (one-sample Wilcoxon signed rank test, z=2.43. p_corr_=0.02) and no significant modulation of the high-β band during execution, or any α/β modulation during observation (all p_corr_=1) (Figure 5B).

The 2×2 repeated measures ANOVA examining the effect of frequency band (α/low-β, high-β) and task (execution, observation) was significant (F_11_=3.36, p=0.006). There was a significant interaction between condition and frequency band (F_1,8_=4.942, p=0.046). Post-hoc contrasts demonstrate greater suppression in α/low-β than high-β during execution (t=3.35, p_corr_=0.02) and no difference between frequency bands during observation (t=0.38, p_corr_=1); comparisons between tasks also demonstrated greater suppression of α/low-β coupling during execution than observation (t=3.15, p_corr_=0.03) and no difference in suppression of high-β (t=0.17, p_corr_=1). Importantly, the PLV measure is independent of power (as long as power is not zero) and changes observed in the power spectrum and PLV dissociated (i.e. suppression of power despite no change in coupling). Therefore these findings cannot be explained by changes in power.

### Motor cortex and GPi alpha/beta-gamma PAC are suppressed during action execution but not action observation

Paired comparisons of dPAC between movement and rest for each recording site and condition across the entire frequency space using non-parametric permutation testing demonstrated significant suppression of coupling between alpha/beta phase and low gamma and HFO amplitude in GPi, and with broadband gamma in motor cortex during movement (Figure 6A/C). In contrast, there was no significant movement-related change in dPAC during action observation in the GPi or motor cortex.

**Figure 6.**
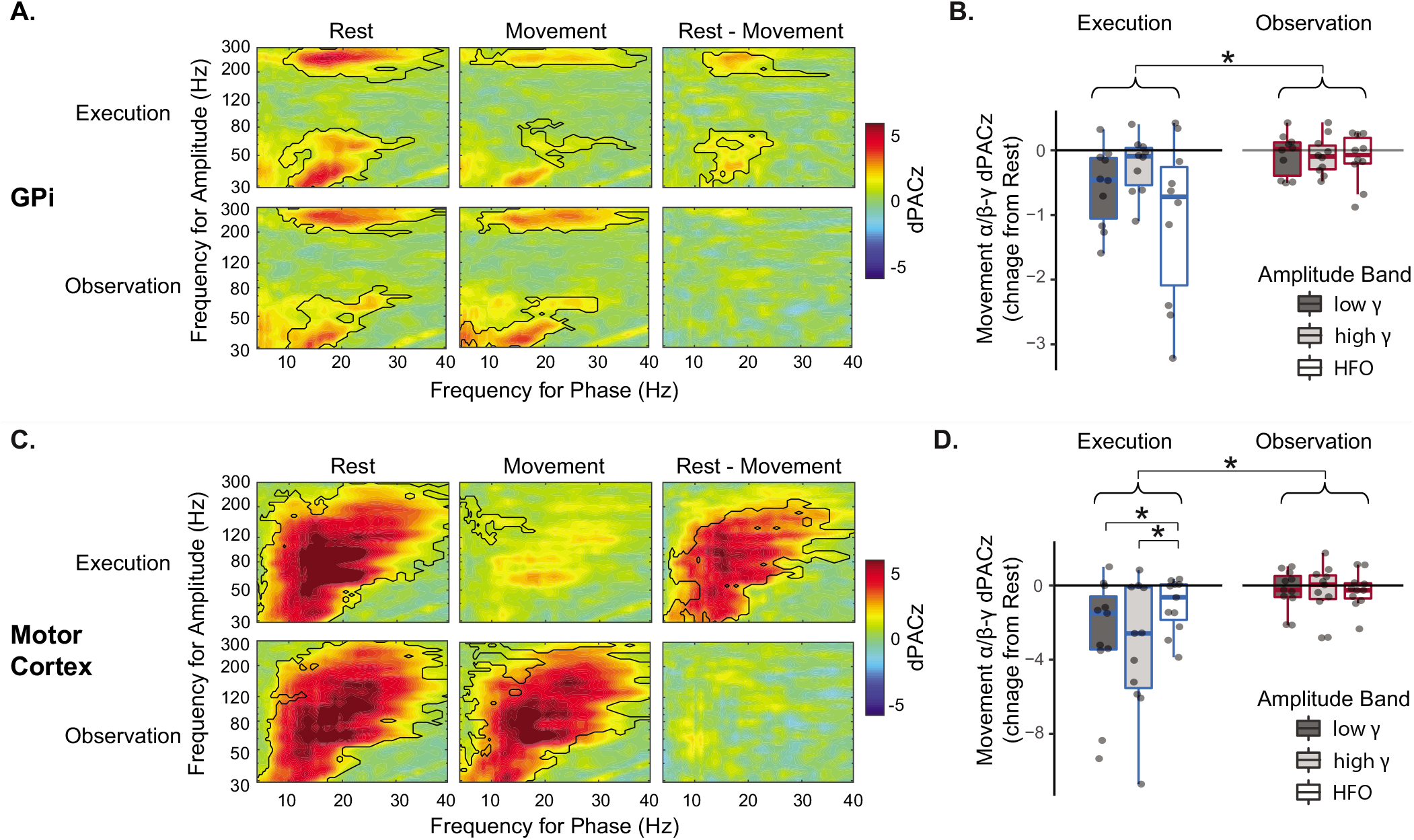
Phase amplitude coupling between alpha/beta phase and gamma/HFO amplitude. dPACz in GPi (A) and motor cortex (C) during movement, rest and their difference demonstrate significant coupling during rest that is suppressed by execution and not by observation. Black outlines indicate significant clusters by non-parametric permutation testing. Boxplots show dPACz modulation in GPi (B) and motor cortex (D) during execution (blue) and observation (red), stratified by amplitude band only as the effect of phase frequency band was not significant for either region. Significant post-hoc contrasts are indicated at top (main effect of task for both regions and trend in task x amplitude band interaction for motor cortex only). (*= p<0.05 post-hoc contrast, Bonferroni corrected)

3-way repeated measures ANOVA examining the effects of phase encoding frequency band (α/low-β, high-β), amplitude modulated frequency band (low-γ, high-γ, HFO) and task (execution, observation) on movement-related modulation were significant in both M1 (F_21_=4.69, p<0.001) and GPi (F_20_=2.27, p=0.004)(Figure 6B/D). In motor cortex, there was a significant main effect of task (F_1,10_=32.9, p<0.001) and trends toward significance in the main effect of amplitude band (F_2,10_=3.19, p=0.056) and the interaction between task and amplitude band (F_2,10_=3.06, p=0.064). Post-hoc contrasts indicate that suppression was greater during execution than observation (t=5.44, p_corr_<0.001) and differences in suppression across amplitude frequency bands was present only during execution (F_2,10_=6.23, p_corr_=0.003) and not during observation (F_2,10_=0.02, p_corr_=0.98); during action execution, suppression was greater in the high-γ and low-γ bands as compared to HFO (t=3.02, p_corr_=0.009 and t=3.09, p_corr_=0.008, respectively). In the GPi, only the main effect of task was significant (F_1,9_=12.69, p<0.001), again due to greater suppression during execution than observation (t=3.56, p<0.001). A trend toward significance was present for the main effect of amplitude modulated frequency band (F_2,9_=3.03, p=0.053) with greater movement suppression for HFO compared to high gamma (t=2.43, p=0.05). Interactions between task and frequency bands were not significant in the GPi. This is likely due to lack of power for the relatively smaller effect sizes, as comodulograms (Figure 6A/C) suggest predominance of suppression in the low-γ and HFO frequencies in the GPi as previously described (Tsiokos et al. 2017; AuYong et al. 2018).

## Discussion

Several previous studies indicate that alpha and beta oscillations in motor cortex and STN are modulated by motor tasks that do not involve overt movement execution (Gastaut and Bert 1954; Pfurtscheller and Neuper 1997; Hari et al. 1998; Kühn et al. 2006a; Marceglia et al. 2009; Alegre et al. 2010; Miller et al. 2010; Brinkman et al. 2014), suggesting that these signal changes may not be directly linked to movement execution. However, previous studies have not examined whether motor tasks also modulate basal ganglia output via the GPi in the absence of overt movement execution. We hypothesized that alpha/beta oscillatory activity in the pallidocortical circuit may be modulated specifically by movement execution, given the GPI’s unique position as the primary output nucleus of the basal ganglia which exerts inhibitory control over the motor system. Our results demonstrate that, in the pallidocortical network, suppression of both local cross frequency coupling and interregional phase coupling occurred only during overt movement, whereas power suppression was a more general property of both overt execution and passive action observation. In addition, we show for the first time that power suppression in alpha/low beta frequencies during action observation extends to the GPi node of the motor system.

### Suppression of local cross frequency and inter-regional phase coupling is specific to movement execution

The dissociation of action execution and observation by pallidocortical phase coupling, but not power modulation, suggests that pallidocortical network interactions are more closely related to movement execution than is power modulation within individual network nodes. Of note, this observation is in contrast to STN power and STN-cortical interactions, which are suppressed across both active and passive motor tasks including action observation (Alegre et al. 2005; Marceglia et al. 2009) and motor imagery (Kühn et al. 2006a; Fischer et al. 2017). We posit that this reflects the unique role of the GPi as the final common output of the basal ganglia, providing a potential mechanism in which gating of actions represented in motor cortex is released through suppression of functional connectivity between GPi and motor cortex.

Similar to cross region coupling, local alpha/beta-gamma PAC in both GPi and motor cortex is suppressed during execution but not during action observation. Previous studies in patients without movement disorders (epilepsy patients) have also demonstrated suppression of motor cortex alpha/beta-gamma PAC with movement (Miller et al. 2012; Yanagisawa et al. 2012), and it is suggested that PAC in the motor cortex may represent a mechanism for controlling movement execution, in which pro-kinetic gamma signals are released from the constraint of alpha/beta phase to allow motor processing (Yanagisawa et al. 2012; de Hemptinne et al. 2013). The excessive PAC observed in PD, may therefore be one mechanism for reduced motor output due to failure of release of pro-kinetic processing (de Hemptinne et al. 2013; de Hemptinne et al. 2015). Our results are in line with this view, given the specificity of PAC suppression to active motor states. Since suppression of alpha/beta pallidocortical phase locking was also seen only during execution, it is possible that the suppression of motor cortex PAC during movement may occur through suppression of GPi-cortical alpha/low-beta functional connectivity, rather than due to suppression of alpha/low-beta power, as reduction of cortical PAC was not seen during observation despite suppression of alpha/low-beta power. The specificity of coupling measures to motor execution in this task, is in line with the view that it may be the heightened network coupling in the β band that contributes to motor symptoms in PD, rather than the degree of local β power.

### Action mirroring in the basal ganglia

Similar activity in neural systems for movement when actions are performed and when they are passively observed has been proposed to provide an efficient mechanism for understanding others’ actions through simulation – observed actions are matched to the observers own motor representation of that action (Rizzolatti and Sinigaglia 2016). Despite a central role of BGTC loops in motor processing and a recent interest in action observation therapies for a variety of motor disorders with prominent basal ganglia dysfunction including PD (Pelosin et al. 2010; Buccino 2014; Caligiore et al. 2017; Pelosin et al. 2018), only few studies have examined whether shared mechanisms for observation and execution of actions extend to subcortical motor regions. Several meta-analyses of neuroimaging studies have suggested that activity during action observation is limited to cortical regions, in contrast to motor execution and motor imagery which also activate subcortical areas (Caspers et al. 2010; Hardwick et al. 2018). However, direct recordings from human STN (Marceglia et al. 2009; Alegre et al. 2010), and now our current data from the GPi, suggest that there may indeed be modulation not detectable by non-invasive neuroimaging methods and that other mechanisms are responsible for differentiating execution and observation.

In this study, we provide the first evidence that spectral power in the GPi is modulated similarly during action observation and execution in parallel to the power modulations previously observed in sensorimotor cortex and STN. Several hypotheses have been put forth regarding the potential role for the basal ganglia during action observation. For example, the basal ganglia may act to reduce or suppress the activity of pyramidal tract neurons to prevent motor output when the motor system is activated by action observation (Bonini 2017), though the suppression of beta power observed here is less consistent with this view given beta suppression is associated with pro-kinetic functions. Alternatively it has been proposed that the basal ganglia may select between multiple potential action representations evoked by an observed action (Caligiore et al. 2013). Finally, as the motor GPi output is likely to be involved in the scaling or vigor of actions (Turner and Anderson 1997; Thura and Cisek 2017), activity in the GPi during action observation may reflect the representation of the dynamics of observed action, which have been shown to be represented in motor cortical beta power modulation (Press et al. 2011). Our work adds to increasing evidence that the basal ganglia, including the GPi, is indeed modulated during action observation (Marceglia et al. 2009; Alegre et al. 2010). Further studies are required to disentangle the above hypotheses regarding specific functional significance during action observation, to understand the potentially differential contributions of GPi and STN to mirroring mechanisms, and to confirm our preliminary data from the visual control task that like the sensorimotor cortex and STN, the GPi modulations observed are specific to observation of actions and not to nonspecific observation of visual stimuli.

### Dissociation between alpha/low-beta and high beta

Studies in Parkinson’s disease have identified functionally distinct bands within the beta frequency range, which classically spans 12-35 Hz oscillations (Oswal et al. 2016; van Wijk et al. 2016; Tsiokos et al. 2017; van Wijk et al. 2017). Our results provide further support for functionally dissociable bands with 2 distinct peaks (around 13 and 23 Hz in this sample) and distributions, which are differentially modulated by execution and observation. Phase coupling modulation by execution was observed only in the alpha/low-beta band and not the high beta band. In addition, power modulation of alpha/low beta was observed during action observation in the absence of high beta modulation, though whether this is attributable to the overall relatively small magnitude of power modulation by action observation or a true band-specific effect remains unclear, as the suppression of power was overall lower in the observation task and no significant frequency band by task interaction was observed.

In previous non-invasive studies of healthy adults, action observation in motor cortex has been shown to be associated with suppression of power in both a lower alpha rhythm (~7-15Hz) and a higher beta rhythm (~15-30 Hz) (Hari 2006; Kilner et al. 2009; Press et al. 2011; Avanzini et al. 2012). It is possible that the relative lack of suppression of the high beta band during action observation in the present study is attributable to PD pathology. Beta synchronization in the BGTC network is excessive in PD, and some studies have proposed high beta synchronization in particular is pathologic (AuYong et al. 2018; Malekmohammadi et al. 2018a). The absence of movement-related suppression of high beta oscillations may be related to network-wide hypersynchrony in this band. However, this is speculative, as the pathological nature of bands is difficult to assess given invasive recordings are limited to clinical populations and others have argued instead that low β synchrony is more closely related to PD pathology (Oswal 2016, Wijk 2016, Litvak 2011). Regardless, clear dissociations between low and high beta bands emerge in our comparison of active and passive motor tasks that warrant further consideration in studies of motor neurophysiology in PD.

### Limitations

Invasive recordings are necessarily driven by clinical indications, which limits the populations available for study and therefore the generalizability to the healthy state as well as the sample size. However, similar movement-related modulation of cortical PAC in healthy populations and patients without movement disorders (Hari et al. 1998; Yanagisawa et al. 2012; Babiloni et al. 2016), and of pallidocortical coupling in patients with dystonia (Tsiokos et al. 2017; van Wijk et al. 2017), suggests that the movement specificity of these coupling measures may well extend beyond the Parkinsonian state. We acknowledge that this work does not exclude the possibility that modulation of phase coupling and phase amplitude coupling in the pallidocortical network may be observed in the absence of motor execution in other tasks (e.g. motor imagery) and future studies to test this hypothesis further are warranted. Nonetheless, the current data demonstrate movement specificity as compared to observation that has not previously been observed in the STN-cortical network.

## Conclusions

We show that coupling in the form of pallidocortical phase locking and local cross frequency coupling in the GPi and motor cortex, but not power modulation, is uniquely modulated by execution suggesting a potentially more direct role of alpha/beta network coupling in motor output. In addition, provide the first evidence of modulation of GPi activity by passive observation of actions in the form of alpha/low beta power suppression.

